# A cullin-RING ubiquitin ligase promotes thermotolerance as part of the Intracellular Pathogen Response in *C. elegans*

**DOI:** 10.1101/586834

**Authors:** Johan Panek, Spencer S. Gang, Kirthi C. Reddy, Robert J. Luallen, Amitkumar Fulzele, Eric J. Bennett, Emily R. Troemel

**Author notes:** Corresponding author: Emily R. Troemel, Address: UC San Diego, 9500 Gilman Dr, #0349, 4205 Bonner Hall, La Jolla, CA 92093-0349. Author Contributions: JP SG KR RL AF EB ET, Designed research JP, SG, EB, ET; Performed research JP, SG, AF; Contributed new reagents or analytic tools SG, RL, KR, ET; Analyzed data JP, AF, SG; Wrote the paper JP, SG, AF, ET.

## Abstract

Intracellular pathogen infection leads to proteotoxic stress in host organisms. Previously we described a physiological program in the nematode *C. elegans* called the Intracellular Pathogen Response (IPR), which promotes resistance to proteotoxic stress and appears to be distinct from canonical proteostasis pathways. The IPR is controlled by PALS-22 and PALS-25, proteins of unknown biochemical function, which regulate expression of genes induced by natural intracellular pathogens. We previously showed that PALS-22 and PALS-25 regulate the mRNA expression of the predicted ubiquitin ligase component cullin *cul-6*, which promotes thermotolerance in *pals-22* mutants. However, it was unclear whether CUL-6 acted alone, or together with other ubiquitin ligase components. Here we use co-immunoprecipitation studies paired with genetic analysis to define the cullin-RING ligase components that act together with CUL-6 to promote thermotolerance. First, we identify a previously uncharacterized RING domain protein in the TRIM family we named RCS-1, which acts as a core component with CUL-6 to promote thermotolerance. Next, we show that the Skp-related proteins SKR-3, SKR-4 and SKR-5 act redundantly to promote thermotolerance with CUL-6. Finally, we screened F-box proteins that co-immunoprecipitate with CUL-6 and find that FBXA-158 promotes thermotolerance. In summary, we have defined the three core components and an F-box adaptor of a cullin-RING ligase complex that promotes thermotolerance as part of the IPR in *C. elegans*, which adds to our understanding of how organisms cope with proteotoxic stress.

**Significance Statement:** Intracellular pathogen infection in the nematode *Caenorhabditis elegans* induces a robust transcriptional response as the host copes with infection. This response program includes several ubiquitin ligase components that are predicted to function in protein quality control. In this study, we show that these infection-induced ubiquitin ligase components form a protein complex that promotes increased tolerance of acute heat stress, an indicator of improved protein homeostasis capacity. These findings show that maintaining protein homeostasis may be a critical component of a multifaceted approach allowing the host to deal with stress caused by intracellular infection.

## Introduction

Maintaining protein homeostasis (proteostasis) after exposure to environmental stressors is critical for organismal survival (1). Several signaling pathways have been identified that help organisms cope with stressors that perturb proteostasis. For example, elevated temperature triggers the conserved Heat Shock Response (HSR) pathway, which helps organisms survive the toxic effects of heat (2). The HSR upregulates expression of chaperones that help with refolding of misfolded proteins, to prevent the formation of protein aggregates and restore proteostasis (1). Disruptions of proteostasis and the formation of protein aggregates in humans are associated with severe neurodegenerative and age-related diseases, such as Alzheimer’s and Huntington’s diseases (1, 3–5).

Pathogen infection can perturb proteostasis and several studies in the nematode *Caenorhabditis elegans* have demonstrated intriguing connections between immune responses to extracellular pathogens and canonical proteostasis pathways (6–11). More recently, examining the *C. elegans* host response to intracellular pathogens has uncovered a novel stress response pathway that promotes proteostasis (12, 13). Microsporidia are intracellular, fungal-like pathogens that are the most common cause of infection of *C. elegans* in the wild, with *Nematocida parisii* being the most commonly found microsporidian species in *C. elegans* (14). *N. parisii* replicates inside the *C. elegans* intestine, and the infection is associated with hallmarks of perturbed proteostasis in the host, such as the formation of large ubiquitin aggregates in the intestine (12). Interestingly, the host transcriptional response to this infection is very similar to the host transcriptional response to another natural intracellular pathogen of the *C. elegans* intestine, the Orsay virus (12, 15, 16). These molecularly distinct pathogens induce a common mRNA expression pattern in *C. elegans* that we termed the “Intracellular Pathogen Response” or IPR (13).

Functional insights into the IPR came from analysis of mutants that constitutively express IPR genes. Forward genetic screens identified two genes encoding proteins of unknown biochemical function called *pals-22* and *pals-25* that comprise an ON/OFF switch for the IPR, with wild-type *pals-25* acting as an activator of the IPR, which is repressed by wild-type *pals-22* (13, 17). Constitutive upregulation of IPR gene expression in *pals-22* loss-of-function mutants is accompanied by a rewiring of *C. elegans* physiology, including increased resistance against natural pathogens like *N. parisii* and the Orsay virus, as well as slowed growth and shortened lifespan. *pals-22* mutants also have increased proteostasis capacity characterized by improved thermotolerance and lowered levels of aggregated proteins (13). All of the *pals-22* mutant phenotypes are reversed in *pals-22 pals-25* loss-of-function double mutants (17). Interestingly, these phenotypes appear to be independent of canonical proteostasis factors (13), such as the transcription factors HSF-1 and DAF-16, which mediate the HSR (18), and the SKN-1/Nrf2 transcription factor, which mediates the proteasomal bounceback response(17, 19). Instead, the proteostasis phenotypes of the IPR require a cullin gene called *cul-6*, which is transcriptionally regulated by *pals-22/pals-25* and by infection (13, 17). Cullins are components of multi-subunit E3 ubiquitin ligases, which are enzymes that catalyze transfer of ubiquitin onto substrate proteins in order to alter their fate (20). Based on these findings we hypothesized that CUL-6-mediated ubiquitylation of target proteins may act as a protein quality control mechanism in the IPR to respond to proteotoxic stress.

CUL-6 belongs to the cullin-RING Ligase (CRL) superfamily, which is found throughout eukaryotes (21). CRLs are multi-subunit enzyme complexes, a subset of which are Skp, cullin, F-box (SCF) complexes consisting of four subunits: a RING-box/RBX protein, a Skp, a cullin and an F-box protein, which serves as a substrate adaptor. Interestingly, the SCF class of ubiquitin ligases appears to have undergone a significant expansion in the evolutionary lineage that gave rise to the nematode *C. elegans* (22, 23). For example, there are around 520 F-box proteins in the *C. elegans* genome (22), in comparison to around 68 in humans (24), 22 in *Drosophila* and 11 in *Saccharomyces cerevisiae* (25). In addition, the number of core SCF components has increased in nematodes, with *C. elegans* having 22 Skp-related proteins in comparison to six in *Drosophila*, just one in *S. cerevisiae*, and one in humans (26). The SCF components upregulated as part of the IPR include not only *cul-6* as mentioned above, but also the Skp-related proteins *skr-3, skr-4 and skr-5,* and several F-box proteins (12). If *cul-6* were functioning as part of a ubiquitin ligase complex to promote proteostasis as part of the IPR, it should be acting with *skrs* and other CRL components. Here we use a combination of biochemistry and genetics to describe how CUL-6 acts together with a RING domain protein, three Skp-related proteins and an F-box protein to promote proteostasis in *C. elegans*.

## Results

### The cullin CUL-6 acts in the intestine and pharynx to promote thermotolerance

In comparison to wild-type animals, *pals-22* loss-of-function mutants have increased thermotolerance, which is reduced to wild-type levels in *pals-22; cul-6* double mutants (13). Our previous results indicated that *pals-22* regulates thermotolerance in the intestine, where it also regulates *cul-6* mRNA expression (13). Because *cul-6* is expressed in the both the intestine and the pharynx (Fig. 1*A*), we investigated where *cul-6* acts to promote thermotolerance. We designed tissue-specific rescue constructs with *cul-6* cDNA using the Mos1-mediated Single Copy Insertion (MosSCI) system (27), to drive expression of GFP-tagged versions of *cul-6* using intestinal (*vha-6p*) or pharyngeal promoters (*myo-2p*). Here we found that both the intestinal and pharyngeal strains expressed GFP::CUL-6 with the expected tissue distribution pattern (Fig. 1*A*). We then crossed these tissue-specific CUL-6 MosSCI transgenes into *pals-22; cul-6* double mutants to test for rescue of thermotolerance. Here we found that expression of *cul-6* in either the intestine or the pharynx was sufficient to increase the thermotolerance of *pals-22; cul-6* double mutants (Fig. 1*B*).

**Figure 1.**
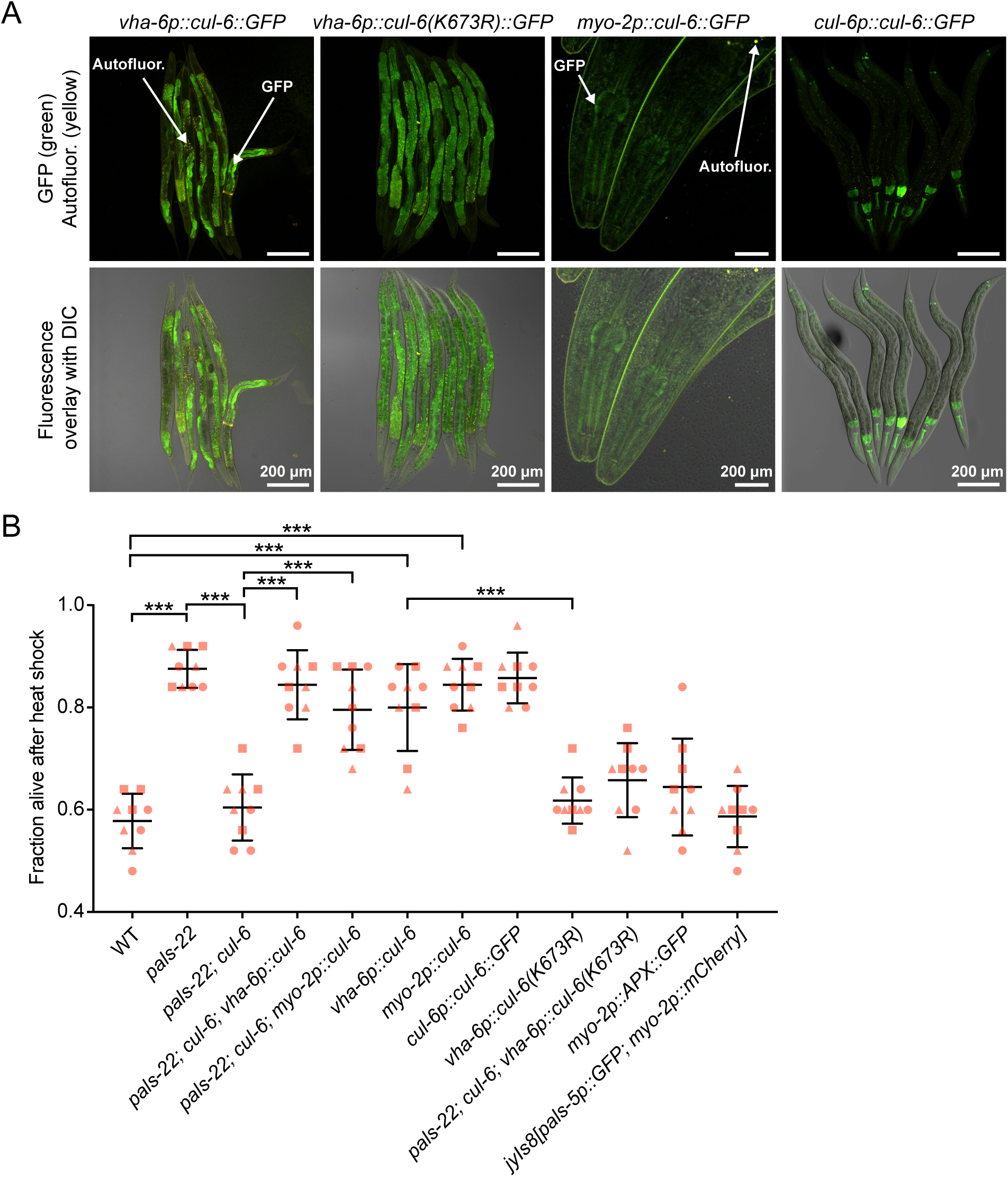
CUL-6 expression in the intestine or in the pharynx promotes thermotolerance. *(A)* Confocal fluorescence images of L4 or adult animals with *cul-6::GFP* transgenes driven by either the *vha-6* or *myo-2* promoter and integrated with MosSCI, or, in the case of *cul-6p::cul-6::GFP*, driven by the endogenous promoter and expressed from a multi-copy array (28). *(B)* Survival of animals after 2 h of 37.5°C heat shock treatment, followed by 24 h at 20°C. Strains were tested in triplicate experiments, with three plates per experiment, 30 animals per plate. The genotypes *myo-2p::APX::GFP* and *jyIs8[pals-5p::GFP; myo-2p::mCherry]* were tested as controls for *myo-2p* driven expression. Each dot represents a plate and different shapes represent the experimental replicates done on different days. Mean fraction alive of the nine replicates is indicated by black bar with errors bars as SD. *** *P* < 0.001, one-way ANOVA with Tukey’s *post-hoc* multiple comparisons test.

Previous studies had only found a functional role for CUL-6 in a *pals-22* mutant background. Here we found that overexpression of CUL-6 from a multi-copy array (CUL-6 tagged at the C-terminus with GFP and 3xFLAG, surrounded by ∼20kb endogenous regulatory region (28)) increased thermotolerance in a wild-type animal background (Fig. 1*A* and *B*). Furthermore, we found that either pharyngeal or intestinal expression of *cul-6* cDNA promoted thermotolerance in a wild type background (Fig. 1*B*). Importantly, transgenic strains with *vha-6* and *myo-2* promoters driving genes other than wild-type *cul-6* did not have increased thermotolerance (see text below, and Fig. 1*B*). Thus, increased expression of *cul-6* in a wild-type background under its own promoter, or only in the intestine or only in the pharynx leads to increased thermotolerance.

We also generated a *myo-3p::cul-6* construct to determine whether expression in body wall muscle could promote thermotolerance. Here, we failed to recover *myo-3p::cul-6* transgenic animals after several rounds of germline injections, suggesting expression of CUL-6 in muscles may be lethal. To quantify this effect, we injected either red fluorescent protein markers together with *myo-3p::cul-6*, or red fluorescent protein markers alone (see Materials and Methods). Here we found none of the eggs expressing red fluorescent protein markers hatched when co-injected with *myo-3p::cul-6* (0/87), while more than half of the eggs hatched when injected with the red fluorescent protein markers alone (48/84). These results suggest that ectopic expression of CUL-6 in muscles is toxic.

The activity of cullin-RING ubiquitin ligases can be increased by neddylation, which is the process of conjugating the ubiquitin-like protein Nedd8 onto a cullin protein at a conserved lysine residue (29). To determine whether CUL-6 might be regulated by neddylation, we mutated the lysine residue that would likely be targeted for neddylation into an arginine, which would be predicted to disrupt neddylation (Fig. S1) (30). We used the MosSCI technique to generate a strain that contains this *vha-6p::cul-6(K673R)* transgene (expression visualized in Fig. 1*A*) and found that it could not rescue the thermotolerance of *pals-22*; *cul-6* mutants (Fig. 1*B*). Furthermore, unlike *vha-6p::cul-6(wild-type)*, the *vha-6p::cul-6(K673R)* transgene did not promote thermotolerance in a wild-type background. These results suggest that CUL-6 requires neddylation to promote thermotolerance, which is consistent with CUL-6 acting as part of a ubiquitin ligase complex.

### Co-immunoprecipitation/mass spectrometry analysis identifies CUL-6 binding partners

Next we performed co-immunoprecipitation mass spectrometry (co-IP/MS) analysis to identify binding partners of CUL-6 (Dataset S1). Here we used the *C. elegans* strain with GFP::3xFLAG-tagged CUL-6, which is functional for thermotolerance (Fig. 1*A*). We also used similar GFP::3xFLAG-tagged strains for PALS-22 and PALS-25 (13, 17). Through analysis of binding partners for PALS-22 and PALS-25, we sought to obtain insight into their biochemical function, which is currently unknown. Two proteins, GFP::3xFLAG alone and an unrelated protein F42A10.5::GFP::3xFLAG, were added as controls for the co-IPs. For each strain used for co-IP/MS analysis we confirmed transgene expression using immunoblotting and microscopy. When we treated animals with the proteasome inhibitor Bortezomib to induce proteotoxic stress and IPR gene expression, we saw an increase in CUL-6 expression by both Western and microscopy analysis, as expected from previous studies (12, 17)(Fig. S2*A* and *B*). CUL-6 expression was seen most strongly in the pharynx and the anterior-most intestinal cells. PALS-22 and PALS-25 were broadly expressed throughout animal and, consistent with previous studies, their expression was not affected by IPR activation (Fig. S2*A* and *B*) (17).

Co-IP/MS of PALS-22 identified 23 binding partners, including PALS-25 as one of the most highly enriched binding partners, as compared to co-IP/MS of control proteins (Fig. S3*A*). PALS-22 also physically associated with PALS-23, which is a PALS protein of unknown function, as well as with F26F2.1, which is a protein of unknown function previously shown to be induced by intracellular infection (12). Co-IP/MS of PALS-25 identified 7 binding partners, with PALS-22 being the most highly enriched hit when compared with co-IP/MS of either control protein (Fig. S3*B*). These reciprocal co-IP results suggest that PALS-22 and PALS-25 are in a physical complex together.

Co-IP/MS of CUL-6 identified 26 significant binding partners. These proteins included predicted SCF ubiquitin ligase components, such as the Skp-related protein SKR-3 and the F-box protein FBXA-158 (Fig. 2*A* and *B*). Additionally, 6 subunits of the 26S proteasome were identified (RPT-3, 4 and RPN-5, 6.1, 8, 9). An SCF ubiquitin ligase complex canonically contains an RBX RING box protein, which interacts with a cullin. *C. elegans* has two RBX proteins, RBX-1 and RBX-2, but neither of these proteins were identified as significant binding partners for CUL-6 in the co-IP/MS. Instead, we identified a single RING domain protein, C28G1.5. Because of results described below, we renamed C28G1.5 as <u>R</u>ING protein acting with <u>cullin</u> and <u>S</u>kr proteins (RCS-1). Interestingly, *rcs-1* mRNA expression, like *cul-6* mRNA expression, is higher in *pals-22* mutants when compared to either wild-type animals (log2 FC=2.81, adjusted p-value= 4.83E-08), or compared to *pals-22 pals-25* mutants (log2 FC=2.10, adjusted p-value=3.898E-07) (17).

**Figure 2.**
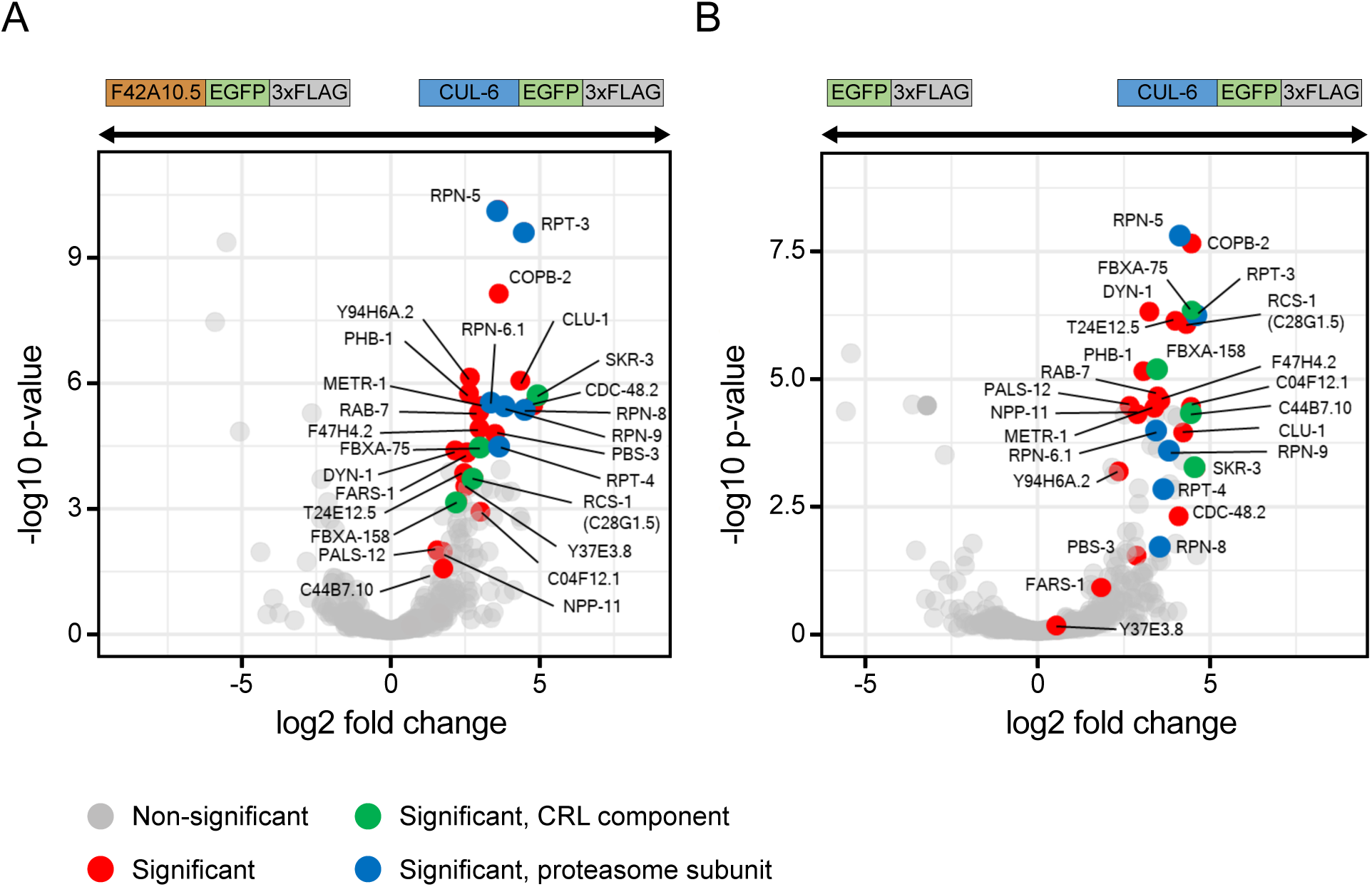
Co-immunoprecipitation mass spectrometry analysis identifies binding partners for CUL-6. Volcano plot of proteins significantly enriched in CUL-6 IP compared to F42A10.5 IP *(A)* or to GFP IP *(B)*. Proteins significantly more abundant compared to either of the control IP’s (GFP alone control or F42A10.5 control, at adjusted *P* < 0.05 and log2 fold change > 1) were considered interacting proteins (Dataset S1). Gray dots indicate non-significant proteins, red dots indicate significant proteins, green dots indicate significant SCF proteins and blue dots indicate significant proteasome subunits.

### RING domain protein RCS-1 acts with CUL-6 to promote thermotolerance in pals-22 mutants

The *rcs-1* gene had no previously described role in *C. elegans* and has two isoforms: *rcs-1b* has a RING domain and a B-box domain, and *rcs-1a* has the RING domain only. The closest potential homolog of *rcs-1* in humans is the tripartite motif-containing protein 23 (TRIM23). The TRIM family is named for having three motifs (RING finger, B box domain, and coiled coil domain), and many TRIM proteins have E3 ubiquitin ligase activity (31). Phylogenetic analysis of the full-length RCS-1 (RCS-1B) indicated that it is part of the *C. elegans* TRIM protein family (Fig. 3*A*) (32), but several *C. elegans* proteins like the ADP-Ribosylation Factor related proteins (ARF-3 and ARF-6) are more closely related to TRIM23 than RCS-1. To determine the expression pattern of RCS-1 we generated transgenic *C. elegans* containing a 3xFLAG and GFP tagged version of RCS-1 as a multi copy array under the control of its endogenous regulatory region (28). The resulting strain expressed GFP throughout the intestine of the worms, with particularly strong expression in the anterior-most intestinal cells (Fig. 3*B*), where CUL-6 is also expressed.

**Figure 3.**
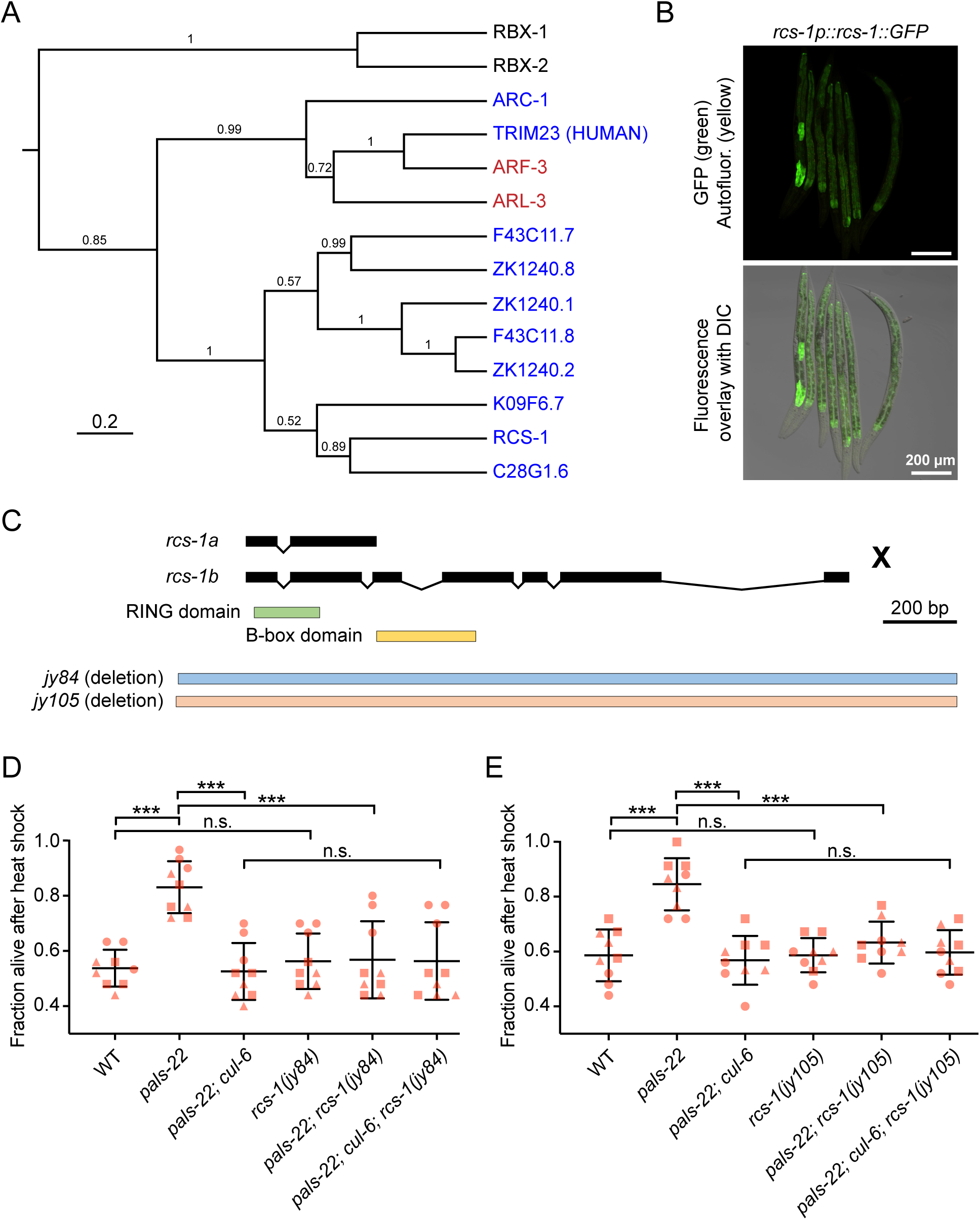
RING domain protein RCS-1 (C28G1.5) promotes thermotolerance in *pals-22* mutants. *(A)* Phylogenetic relationships of RCS-1 protein with TRIM23 homologs proteins (in red), canonical RBX proteins (in black) and known TRIM proteins (in blue – all are *C. elegans* except noted human gene). The tree was built from a protein alignment using the Bayesian MCMC method. Posterior probabilities are indicated on the branches. *(B)* Confocal fluorescence images of L4 animals with *rcs-1::GFP* driven by the endogenous promoter and expressed from a multi-copy array (28). *(C) rcs-1* isoforms and exon/intron structures. Protein domains are colored in green and yellow. *jy84* and *jy105* are deletion alleles. *(D* and *E)* Survival of animals after 2 h of 37.5°C heat shock treatment, followed by 24 h at 20°C. Strains were tested in triplicate experiments, with three plates per experiment, 30 animals per plate. Each dot represents a plate, and different shapes represent the experimental replicates done on different days. Mean fraction alive of the nine replicates is indicated by black bar with errors bars as SD. *** *P* < 0.001, one-way ANOVA with Tukey’s *post-hoc* multiple comparisons test.

Next we used CRISPR-Cas9 to generate two independent deletion alleles of *rcs-1* (*jy84* and *jy105*), which we crossed into *pals-22* mutants (Fig. 3*C*). We found that, for both *rcs-1* alleles, *pals-22; rcs-1* double mutants had thermotolerance similar to wild-type animals, indicating that *rcs-1* is required for the increased thermotolerance of *pals-22* mutants (Fig. 3*D* and *E*). Similar to *cul-6*, *rcs-1* mutations had no effect in a wild-type background (Fig. 3*D* and *E*). If RCS-1 were acting in a SCF together with CUL-6, then loss of *rcs-1* would not further lower thermotolerance in a *pals-22; cul-6* mutants. Indeed, we found that *pals-22; cul-6; rcs-1* triple mutants had a similar level of thermotolerance to *pals-22; cul-6* mutants, as well as to *pals-22; rcs-1* mutants and wild-type animals. These findings, together with CUL-6 co-IP results and RCS-1::GFP expression pattern, are consistent with RCS-1 being the RING domain protein that acts with CUL-6 in a ubiquitin ligase complex.

### SKP-related proteins SKR-3, SKR-4 and SKR-5 act redundantly to promote thermotolerance in pals-22 mutants

In addition to a cullin and a RING protein, SCF ubiquitin ligase complexes contain a Skp protein. Expression of three Skp-related genes (*skr-3, skr-4 and skr-5*) is upregulated by both intracellular infection and mutation of *pals-22*, similar to *cul-6* expression (12, 13). We previously found that mutation of either *skr-3, skr-4* or *skr-5* alone in a *pals-22* mutant background had no effect on thermotolerance (Fig. 4*A*) (13). Therefore, we made all of the possible combinations of *skr-3*, *skr-4* and *skr-5* as double mutants and then crossed them into a *pals-22* mutant background to determine whether they may act redundantly. Here we found that *pals-22; skr-3 skr-5* and *pals-22; skr-5 skr-4* triple mutants had a significant reduction in thermotolerance compared to *pals-22* mutants, with levels similar to wild-type animals. In contrast, *pals-22; skr-3 skr-4* mutants had thermotolerance similar to *pals-22* mutants (Fig. 4*B*). These results indicate that either SKR-3, SKR-4, or SKR-5 can act together with CUL-6 in a SCF to promote thermotolerance, with SKR-5 being the most important. Consistent with the idea that SKR-5 acts together with CUL-6 and RCS-1, we found that SKR-5::GFP::3xFLAG under control of endogenous regulatory regions is strongly expressed in the anterior-most cells of the intestine (Fig. 4*C*).

**Figure 4.**
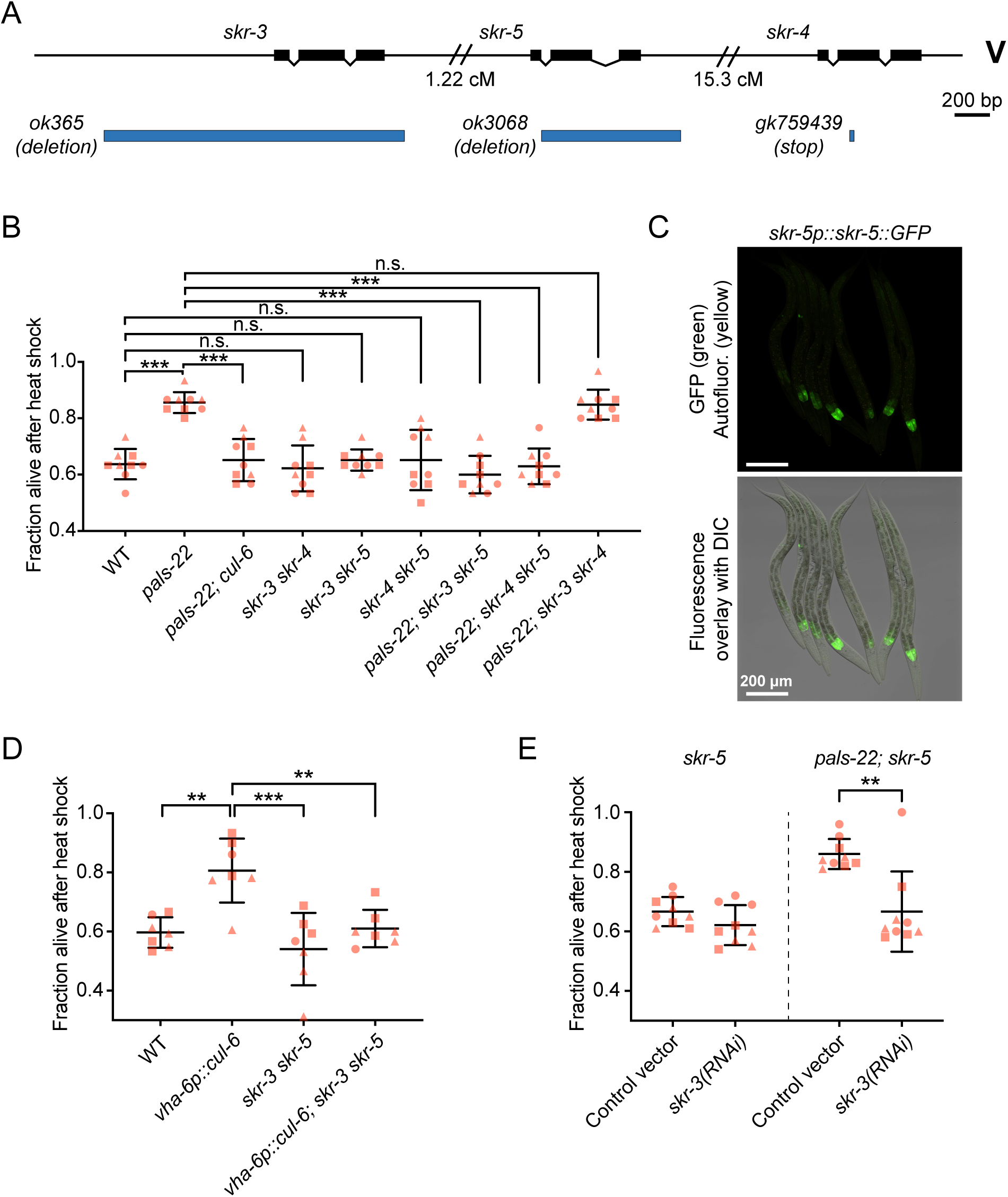
SKR-3, SKR-4 and SKR-5 act redundantly to promote thermotolerance in *pals-22* mutants. *(A) skr-3, skr-4 and skr-5* gene exon/intron structure. *ok365* and *ok3068* are deletion alleles, *gk759439* is a premature stop mutation. *(B)* Survival of animals after 2 h of 37.5°C heat shock treatment, followed by 24 h at 20°C. *** *P* < 0.001, one-way ANOVA with Tukey’s *post-hoc* multiple comparisons test. n = 9 replicates per strain. *(C)* Confocal fluorescence images of L4 animals with *skr-5::GFP* driven by the endogenous promoter and expressed from a multi-copy array (28). *(D)* Survival of animals after 2 h of 37.5°C heat shock treatment, followed by 24 h at 20°C. *** *P* < 0.001, ** *P* < 0.01, one-way ANOVA with Tukey’s *post-hoc* multiple comparisons test. n = 7 replicates per strain. *(E)* Survival of animals after 2 h of 37.5°C heat shock treatment, followed by 24 h at 20°C. *skr-5* and *pals-22*; *skr-5* mutants were fed on (R)OP50 expressing either L4440 (control vector) or *skr-3* RNAi. *** *P* < 0.001, two-way ANOVA with Sidak’s multiple comparisons test. n = 9 replicates per condition. For *(B, D* and *E*) strains were tested in triplicate experiments, 30 animals per plate. Mean fraction alive of the replicates is indicated by black bar with errors bars as SD. Each dot represents a plate, and different shapes represent the experimental replicates done on different days.

Our results indicated than CUL-6 overexpression in the intestine was sufficient to increase thermotolerance in a wild-type background (Fig. 1*B*). To investigate whether CUL-6 acts together with SKR proteins in this context, we crossed the *skr-3 skr-5* double mutant into the CUL-6 overexpressing strain (*vha-6p::cul-6*). As predicted, the resulting strain had a thermotolerance phenotype comparable to wild-type animals, consistent with CUL-6 acting together with SKR proteins (Fig. 4*D*).

Next we sought to use RNA interference (RNAi) to further validate the redundancy of SKR proteins acting with CUL-6. However, we found that *pals-22* mutants do not have increased thermotolerance compared to wild-type animals when fed on the standard *E. coli* bacteria used for feeding RNAi (HT115) (Fig. S4*A*). This effect may be due to dietary differences between HT115 and the OP50 strain used for thermotolerance experiments described above (33). Therefore, we tested thermotolerance with an OP50 strain (R)OP50 that was modified to enable feeding RNAi studies (34). Here we found that *pals-22* mutant animals fed on these RNAi-competent OP50 bacteria (transformed with empty vector L4440) have increased thermotolerance compared to wild-type animals, which is restored back to wild-type levels in *pals-22; cul-6* mutants (Figure S4*A*). Using this system, we successfully knocked down expression of *cul-6* with RNAi as assessed by lowered CUL-6::GFP::3xFLAG transgene expression (Figure S4*B* and *C*). Here we found that *cul-6* RNAi suppressed the enhanced thermotolerance of *pals-22* mutants (Figure S4*D*). With this system, we then confirmed that *skr-3* and *skr-5* act redundantly to promote thermotolerance, as RNAi against *skr-3* suppressed thermotolerance of *pals-22; skr-5* double mutants but not in *skr-5* mutants (Fig. 4*E*).

### Analysis of CUL-6, SKR-3,4,5 and RCS-1 in other phenotypes mediated by pals-22

Previous analysis of IPR genes indicated that RNAi knock-down of *cul-6*, *skr-3* or *skr-5* increased susceptibility to intracellular infection in a sterile but otherwise wild-type background *C. elegans* strain (12). Here we investigated whether *cul-6* mutants, *skr-3 skr-5* or *skr-4 skr-5* double mutants, or *rcs-1* mutants could suppress the increased pathogen resistance of *pals-22* mutants to microsporidia. In contrast to the complete suppression we found for increased thermotolerance of *pals-22* mutants, we found only a minor suppression of pathogen resistance by these mutants in a *pals-22* mutant background (Fig. S5). We also found that mutations in *rcs-1*, *skr-3 skr-4*, *skr-3 skr-5,* or *skr-5 skr-4* did not suppress the slowed developmental rate in *pals-22* mutants, similar to *cul-6* (Fig. S6). Therefore, *cul-6, rcs-1,* and *skr-3,4,5* appear to be important for executing the thermotolerance phenotype of *pals-22* mutants, but not other phenotypes.

### FBXA-158 acts with CUL-6 to promote thermotolerance in pals-22 mutants

Next we sought to identify the F-Box protein that functions as a substrate adaptor with CUL-6, RCS-1 and SKR-3,4,5. Here we used RNAi to screen through the F-box proteins that were found to physically associate with CUL-6 from co-IP/MS. In addition to FBXA-158 and FBXA-75 as significant hits, there were other F-box proteins that we identified below the significance threshold, including FBXA-54, FBXA-188 and FBXA-11. Because there were RNAi clones available for *fbxa-158*, *fbxa-54*, *fbxa-188* and *fbxa-11* we used the (R)OP50 RNAi system to screen these genes in a *pals-22* mutant background. When treated with RNAi, only *fbxa-158* showed significant suppression of the increased thermotolerance phenotype (Fig. 5*A*). To further validate the results from our initial screen we also tested *fbxa-158* RNAi in the CUL-6 overexpression strain *vha-6p::cul-6*. Here, *fbxa-158* RNAi treatment similarly suppressed the *vha-6p::cul-6* increased thermotolerance phenotype (Fig. 5*B*). Moreover, when treated with *fbxa-158* RNAi, wild-type animals did not show further decreased thermotolerance, thus indicating that FBXA-158 is not acting independently from CUL-6 (Fig. 5*B*).

**Figure 5.**
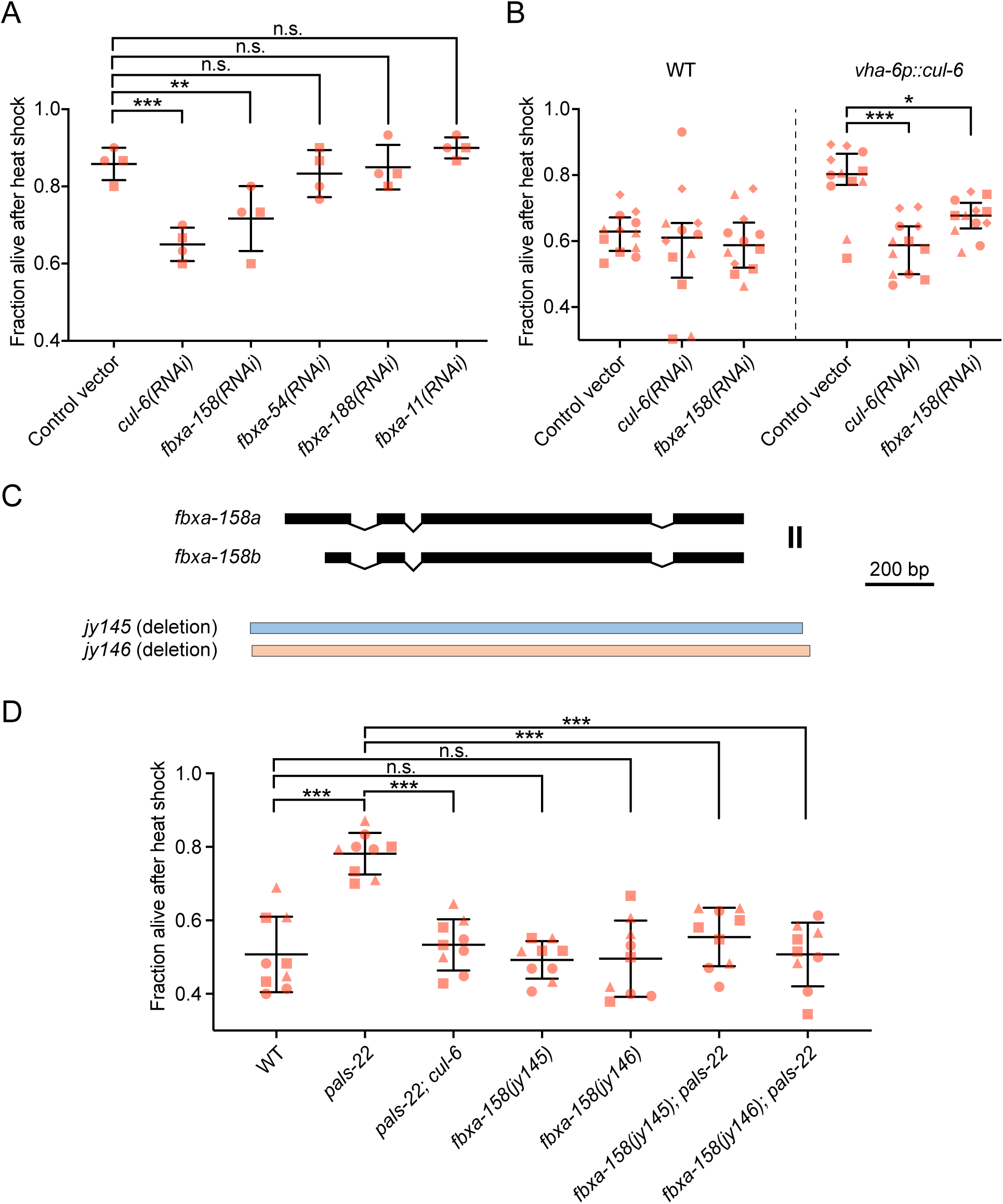
FBXA-158 promotes thermotolerance in *pals-22* mutants. *(A)* Survival of animals after 2 h of 37.5°C heat shock treatment, followed by 24 h at 20°C. *pals-22* mutants were fed on (R)OP50 expressing either L4440 (control vector) or RNAi for the indicated genes. *pals-22* mutants were tested in duplicate experiments, with two plates per experiment, 30 animals per plate. *** *P* < 0.001, ** *P* < 0.01, one-way ANOVA with Tukey’s *post-hoc* multiple comparisons test. *(B)* Survival of animals after 2 h of 37.5°C heat shock treatment, followed by 24 h at 20°C. Wild type and *vha-6p::cul-6* animals were fed on (R)OP50 expressing either L4440 (control vector), *cul-6* or *fbxa-158* RNAi. Strains were tested in quadruplicate experiments, with three plates per experiment, 30 animals per plate. *** *P* < 0.001, * *P* < 0.05, two-way ANOVA with Sidak’s multiple comparisons test. *(C) fbxa-158* isoforms and exon/intron structures. *jy145* and *jy146* are deletion alleles. *(D)* Survival of animals after 2 h of 37.5°C heat shock treatment, followed by 24 h at 20°C. *** *P* < 0.001, one-way ANOVA with Tukey’s *post-hoc* multiple comparisons test. *(A, B* and *D)* each dot represents a plate, and different shapes represent the experimental replicates done on different days. Mean fraction alive of the replicates is indicated by black bar with errors bars as SD.

To confirm our RNAi results we used CRISPR-Cas9 to generate two independent deletion alleles of *fbxa-158* (*jy145* and *jy146*) and crossed them with *pals-22* mutants (Fig. 5*C*). When tested for thermotolerance, both *fbxa-158* deletion alleles suppressed the increased thermotolerance of *pals-22* mutants while in wild-type animals the survival rate was unchanged (Fig. 5*D*). Similar to *cul-6, rcs-1* and *skr-3,4,5*, *fbxa-158* mRNA expression is higher in *pals-22* mutants when compared to wild-type animals (log2 FC=2.81, adjusted p-value= 4.83E-08), and in *pals-22* mutants compared to *pals-22 pals-25* mutants (log2 FC=6.95, adjusted p-value=1.2349E-06) (17). *fbxa-158* expression is also enriched in the intestine (35). Together, these results indicate that FBXA-158 acts as an F-box adaptor protein in a CUL-6/RCS-1/SKR-3,4,5 ubiquitin ligase complex that promotes thermotolerance in *C. elegans*.

## Discussion

Thermal stress is one of many types of proteotoxic stress that can impair organismal health and survival. Here we used a combination of genetics and biochemistry to broaden our understanding of a recently identified proteostasis pathway called the IPR, which enables animals to survive exposure to thermal stress in a manner distinct from the canonical heat shock response (HSR). Specifically, we demonstrate that overexpression of CUL-6/cullin alone promotes thermotolerance, and it can act in either the intestine or the pharynx of *C. elegans*. Importantly, we found that CUL-6 acts together with other ubiquitin ligase components, including a previously uncharacterized RING protein we named RCS-1, as well as the Skp-related proteins SKR-3, SKR-4, and SKR-5 and the F-box protein FBXA-158 (Fig. 6). We propose that this RCS-1/CUL-6/SKR-3,4,5/FBXA-158 ubiquitin ligase complex is able to target proteins for ubiquitin-mediated proteasomal degradation and that this activity is a critical part of the IPR program. Consistent with this model, our co-IP/MS identified several proteasomal subunits that interact with CUL-6 (Fig. 2).

**Figure 6.**
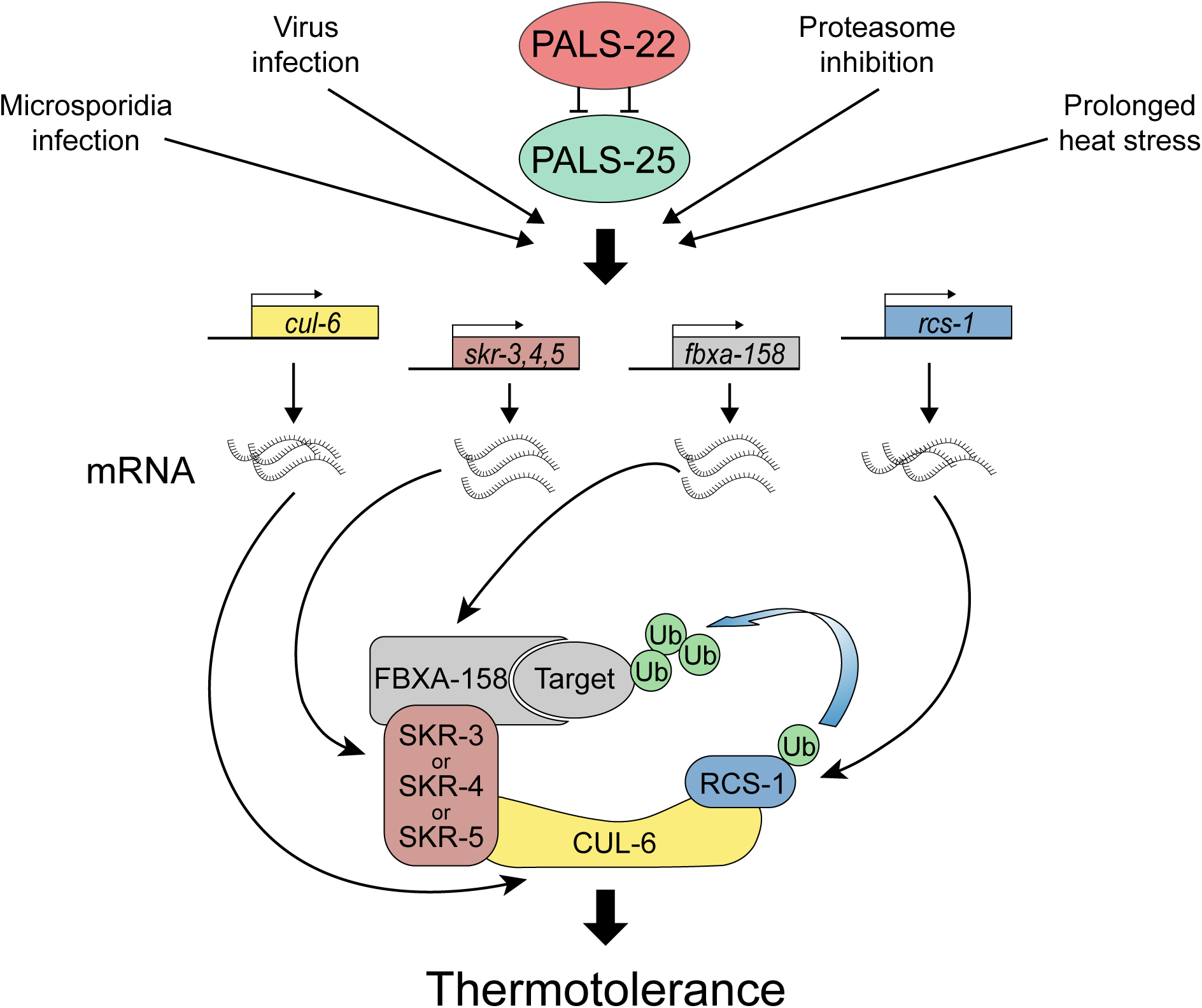
Model for a RCS-1/CUL-6/SKR/FBXA-158 ubiquitin ligase that promotes proteostasis.

We also investigated protein-protein interactions of the PALS-22 and PALS-25 proteins, which comprise an ON/OFF switch in the IPR that regulates mRNA expression of *cul-6, skr-3, skr-4, skr-5, rcs-1* and *fbxa-158*. Previous studies indicated that *pals-22* and *pals-25* are in the same operon and interact genetically, and our co-IP/MS studies indicate they also interact biochemically. Notably, the *pals* gene family has expanded in the *C. elegans* genome, with 39 genes in *C. elegans*, in comparison to only 1 *pals* gene each in mouse and human (36). Although the divergent ‘*pals’* protein signature that defines PALS proteins is of unknown function, PALS-22 does have weak homology with F-box proteins (36), leading to the speculative idea that PALS proteins function as adaptor proteins in ubiquitin ligase complexes.

Like the *pals* gene family, *C. elegans* has a greatly expanded repertoire of SCF ligase components. This expansion has been suggested to reflect the results of an arms race against intracellular pathogens, as SCF components are among the most rapidly diversifying genes in the *C. elegans* genome (23). The most dramatically expanded class of SCF components includes ∼520 F-box adaptor proteins in *C. elegans* (22). *C. elegans* also has 22 SKR proteins, in comparison to only 1 Skp in humans, indicating there has been an expansion of core SCF components as well. Here we identified SKR-3 as a binding partner for CUL-6, which is consistent with previous 2-hybrid results (26). Interestingly, we found redundancy in the role of SKRs at the phenotypic level: either SKR-3, SKR-4 or SKR-5 appear capable of acting with CUL-6 to promote thermotolerance, with SKR-5 being the most important. We also found that FBXA-158, one of the two F-box proteins identified as binding partners of CUL-6 by co-IP/MS acts to promote thermotolerance. Together, with the Skp-related proteins, the F-box protein are responsible for substrate specificity of the SCF (37). Hence the identification of FBXA-158 and SKR-3,4,5 as member of the complex will help in future studies to identify what specific proteins are ubiquitylated by a CUL-6 SCF.

Canonical CRLs contain an RBX protein as the RING domain protein, which interacts with both a cullin and an E2 ubiquitin ligase (21). However, our co-IP/MS studies with CUL-6 did not identify RBX-1 or RBX-2 as interacting partners for CUL-6, but rather identified RCS-1. Given our genetic and biochemical results, we propose that RCS-1 plays the same role as an RBX protein would in a canonical SCF complex (Fig. 6). RCS-1 does not have a clear human ortholog, but the closest human protein is TRIM23 (38). Interestingly, the TRIM family contains 68 genes in humans, many of which encode single subunit E3 ubiquitin ligases, including those that restrict viral infection, and regulate inflammatory signaling (31). *C. elegans* has 18 TRIM proteins, and they appear to have a simpler structure than human TRIM proteins, given that absence of additional motifs in the C-terminal domains normally found in the majority of mammalian TRIM proteins (32). If these other *C. elegans* TRIM proteins can act in SCF ligases like RCS-1, it suggests there may also be an expansion of the RING core SCF components in *C. elegans*, in addition to the previously described expansion of SKRs and adaptor proteins.

The role of CUL-6 in promoting thermotolerance was first demonstrated in *pals-22* mutants, where there is an upregulation of *cul-6* mRNA as well as several other SCF components, including *skr-3, skr-4, skr-5, rcs-1* and *fbxa-158* (13, 17). However, here we found that animals overexpressing only CUL-6, without overexpression of the other components of the SCF, have increased thermotolerance. One explanation for this result is that CUL-6 is the limiting factor in a SCF that promotes thermotolerance in the IPR. Consistent with this idea, other components of the complex like SKR-3,4,5 are functionally redundant for thermotolerance, so even basal expression level might be sufficient to build a functional SCF, once CUL-6 expression increases past a certain threshold level.

What substrate(s) is targeted by the RCS-1/CUL-6/SKR-3,4,5/FBXA-158 ubiquitin ligase complex? It is possible that the effects of this complex are mediated through targeting a single regulatory protein for ubiquitylation and degradation. For example, ubiquitylation of the DAF-2 insulin receptor by the ubiquitin ligase CHIP can alter DAF-2 trafficking, and it appears that ubiquitylation of just this one target has significant effects on proteostasis in *C. elegans* (39). In contrast, CRL2 and CRL4 complexes in humans have recently been shown to target the C-termini of a large number of proteins for degradation as part of a newly identified protein quality control system (40, 41), Given the modular nature of the SCF ligase family, the large number of F-box substrate adaptor proteins in *C. elegans*, and the redundancy we found in the SKR-3,4,5 proteins, it seems possible that there are multiple substrates and adaptors used in the IPR. An exciting possibility is that the IPR involves distinct CRLs that ubiquitylate several different targets to improve proteostasis and tolerance against environmental stressors, including infections. Identifying these factors will be the subject of future studies.

## Materials and Methods

### Cloning and generation of cul-6 tissue-specific rescue strains

A full list of strains used in this study is in Table S1. A full list of constructs used in this study is in Table S2. To generate the *vha-6p::CUL-6* transgene (pET499), *vha-6p::SBP::3XFLAG*, *cul-6* cDNA, and the *unc-54* 3’ UTR were assembled in pCFJ150 using Gateway LR. To generate *myo-2p::CUL-6* (pET686) and *myo-3p::CUL-6* (pET687) constructs, the promoters *myo-2p*, *myo-3p* and the pET499 linearized backbone without the *vha-6p* promoter were amplified by PCR from pCFJ90, pCFJ104 and pET499, respectively and assembled by Gibson recombination (42). To generate the *cul-6(K673R)* construct (pET688) a 86 bp single strand oligonucleotide and a linearized pET499 backbone that had been amplified by PCR were assembled by Gibson recombination. To generate the *spp-5p::3XFLAG::GFP* (pET555) transgene, *spp-5p::3XFLAG::GFP* and *let-858* 3’ UTR were assembled in pCFJ150 using Gateway LR. Mos1-mediated Single Copy Insertion (MosSCI) was performed as described previously (27). Briefly, the plasmid of interest (25 ng/µl) was injected with pCFJ601 (50 ng/µl), pMA122 (10 ng/µl), pGH8 (10 ng/µl), pCFJ90 (2.5 ng/µl), and pCFJ104 (5 ng/µl) into the EG6699 strain. Injected animals were incubated at 25°C for 7 days and then subjected to heat shock for 2h at 34°C. After 5 h non-Unc animals were selected and the presence of the insertion was verified by PCR and sequencing.

Transgenic strains with TransgeneOme fosmids (Table S2) were generated as extrachromosomal arrays (28) by injecting into *ttTi5605; unc-119(ed3)* worms (strain EG6699) and then selecting non-Unc worms.

### Lethality scoring of myo-3p::cul-6 tissue-specific rescue strains

To score the lethality of *myo-3p::cul-6* expression, worms were injected with a complete MosSCI mix (see above) containing either *myo-3p::cul-6* as the plasmid of interest, or water. Injected animals were incubated at 25°C and after 1 day, eggs expressing red fluorescence were transferred onto new plate at 25°C for 24 h. The hatching ratio of transferred eggs was then scored for both conditions. The assay was repeated two independent times.

### Thermotolerance assays

Animals were grown on standard NGM plates at 20°C. L4 stage animals were transferred onto fresh NGM plates seeded with OP50-1 and then subjected to heat shock for 2 h at 37.5°C. The plates were then placed in a single layer on a benchtop at room temperature for 30 min, and then transferred to a 20°C incubator. Then, 24 h later the survival was scored in a blinded manner. Worms not responding to touch were scored as dead, and 30 worms were scored per plate. Three replicate plates were scored for each strain per experiment, and each experiment was performed at least three independent times. Statistical significance was tested using one-way ANOVA and Tukey’s HSD for *post-hoc* multiple comparisons.

### Co-immunoprecipitation

Each sample for co-IP/MS was prepared in 3 independent experimental replicates. For each sample, 200,000 synchronized L1 animals were transferred onto NGM plates and grown for 48 h at 20°C. Bortezomib was added to reach a final concentration of 22 µM or the equivalent volume of DMSO for the control plates. After 6 h at 20°C worms were washed off of the plates with M9, washed twice with M9, resuspended in 500 µl of ice-cold lysis buffer (50mM HEPES, pH7.4, 1mM EGTA, 1mM MgCl2, 100mM KCl, 1% glycerol, 0.05% NP40, 0.5mM DTT, 1x protease inhibitor tablet) and immediately frozen dropwise in liquid N_2_. Frozen pellets were ground into powder with a pre-chilled mortar and pestle. Protein extracts were spun for 15 min 21,000g at 4°C and supernatants were filtered on 0.45 µm filters (Whatman). Protein concentration was determined using Pierce 660nm protein assay and adjusted to 1 µg/µl with fresh lysis buffer. 1 mg of each sample was mixed with 25 µl of ANTI-FLAG M2 Affinity Gel (Sigma) and incubated at 4°C with rotation (12 rpm) for 1h. The resin was washed twice with 1 ml lysis buffer, twice with 1 ml lysis buffer for 5min, twice with 1 ml wash buffer (50mM HEPES, pH7.4, 1mM MgCl2, 100mM KCl) and 20min with 1 ml wash buffer with rotation. The liquid was removed and the beads were then stored at -80°C.

### Trypsin digestion

The immunoprecipitated proteins bound to the beads were digested overnight in 400 ng trypsin (Sigma, V511A) in 25 mM ammonium bicarbonate (Sigma) at 37°C. Samples were then reduced with 1mM final concentration of Dithiothreitol (DTT, Acros Organics) for 30min and alkylated with 5mM final concentration of Iodoacetamide (IAA, MP Biomedicals, LLC) for 30min in dark. The peptides were extracted from the beads by adding 50 µL of 5% Formic acid (Sigma). The extraction was repeated one more time and the eluted peptides were combined. Digested peptides were desalted using Stage-Tip, C18 peptide cleanup method. The eluates were vacuum dried, and peptides were reconstituted in 15 µL of 5% Formic acid, 5% Acetonitrile solution for LC-MS-MS analysis.

### LC-MS-MS Analysis

Samples were analyzed in triplicate by LC-MS-MS using an EASY-nLC 1000 HPLC (Thermo Scientific) and Q-Exactive mass spectrometer (Thermo Scientific, San Jose, CA) as described previously (43) with the following modifications. The peptides were eluted using a 60min acetonitrile gradient (45 min 2%-30% ACN gradient, followed by 5 min 30-60% ACN gradient, a 2min 60-95% ACN gradient, and a final 8min isocratic column equilibration step at 0% ACN) at 250nL/minute flow rate. All the gradient mobile phases contained 0.1% formic acid. The data dependent analysis (DDA) was done using top 10 method with a positive polarity, scan range 400-1800 m/z, 70,000 resolution, and an AGC target of 3e6. A dynamic exclusion time of 20 s was implemented and unassigned, singly charged and charge states above 6 were excluded for the data dependent MS/MS scans. The MS2 scans were triggered with a minimum AGC target threshold of 1e5 and with maximum injection time of 60 ms. The peptides were fragmented using the normalized collision energy (NCE) setting of 25. Apex trigger and peptide match settings were disabled.

### Peptide and Protein Identification and Quantification

The RAW files obtained from the MS instrument were converted into mzXML format. The SEQUEST search algorithm was used to search MS/MS spectra against the concatenated target decoy database comprised of forward and reverse, reviewed *C. elegans* FASTA sequences from Uniprot (downloaded on 6/8/2015) along with GFP and *E. coli* sequences appended in the same file. The search parameters used were as follows: 20 ppm peptide mass tolerance; 0.01 Da fragment ion tolerance; Trypsin (1 1 KR P) was set as the enzyme; maximum 2 missed cleavages were allowed; Oxidation on methionine (15.99491 Da) and n-term acetylation (42.01056 Da) were set as differential modifications; static modification (57.02146 Da) was set on cysteine for alkyl modification. Peptide matches were filtered to a peptide false discovery rate (FDR) of 2% using the linear discrimination analysis. The protein level matches were filtered at 2% FDR using the protein sieve analysis. The spectral counts from the triplicates were then summed and used for the data analysis.

### Analysis of mass spectrometry data

Peptide spectral counts were used to calculate fold change ratio, p-value and adjusted p-value between sample IPs and control IPs (GFP and F42A10.5) using the DEP package in R (44). Briefly the data were filtered to keep only the peptides present in at least two replicates in one condition. Filtered data were normalized using Variance Stabilizing Normalization. Missing values were imputed using the MiniProb method from the DEP package by randomly selecting values from a Gaussian distribution centered on a minimal value of the dataset. Fold change ratio and adjusted p-values were calculated. Proteins with adjusted *P* < 0.05 and log2 fold change > 1 in comparison with at least one of the controls were considered as significant. Any protein with a negative log2 fold change with one control or the other (*i.e.* more affinity to the control protein than the tested bait) was considered as non-significant.

### Phylogenic analysis of RCS-1

Amino acid sequences of 15 proteins were aligned using MUSCLE (version 3.7) and trimmed with trimAI (version 1.3) (45) using phylemon2 online platform (46). Bayesian Markov chain Monte Carlo inference (LG + I + G + F) was performed using BEAST (version 1.10.4) (47). Analysis was run using a Yule model tree prior, an uncorrelated relaxed clock (lognormal distribution) and a chain length of 10 million states sampled every 1,000 iterations. Results were assessed with Tracer (version 1.7.1), maximum clade credibility tree was built after a 25% burn-in. Posterior probability values greater than 0.5 are marked on branch labels.

### CRISPR deletions of rcs-1 and fbxa-158

To generate deletions of *rcs-1* and *fbxa-158,* a co-CRISPR strategy was used, adapted from the IDT proposed method for *C. elegans*. To generate the ∼2.18 kb deletion alleles of *rcs-1* we designed two crRNA encompassing the whole *rcs-1* locus (crRNA1: 5’-GTTTGTTGAAGGAAATGCACAGG-3’, crRNA2: 5’-GGTTTCCTATAGCTGTGACACGG-3’). These two oligonucleotides were synthetized by IDT and used with a crRNA targeting the *dpy-10* gene (*dpy-10* crRNA3: 5’-GCTACCATAGGCACCACGAG-3’) and assembled with commercial tracrRNA (50 µM crRNA1 and crRNA2, 25 µM *dpy-10* crRNA, and 40 µM tracrRNA). After an annealing step for 5 min at 95°C, the resulting guide RNA mixture was added to CAS9-NLS protein (27 µM final – ordered from QB3 Berkeley) and microinjected into N2 worms. F1 Dpy progeny were screened by PCR, confirmed by sequencing for a deletion and positive lines were backcrossed 4 times to the N2 background before testing. The same co-CRISPR strategy was used to generate the ∼1.65 kb deletion alleles of *fbxa-158*. Two crRNAs encompassing the whole *fbxa-158* locus (crRNA1: 5’-ATAGTCGGGTACAAAACAAATGG-3’, crRNA2: CTACTCCATCTTTAAGAACACGG-3’) were used with the same *dpy-10* crRNA and assembled as described above. Deletion positive lines were backcrossed 2 times to the N2 background before testing.

### RNA interference assays

RNA interference assays were performed using the feeding method. Overnight cultures of HT115 or OP50 strain (R)OP50 modified to enable RNAi (34) (gift from Meng Wang lab, Baylor College of Medicine) were plated on RNAi plates (NGM plates supplemented with 5 mM IPTG, 1 mM carbenicillin) and incubated at 20°C for 3 days. Gravid adults were transferred to these plates, and their F1 progeny (L4 stage) were transferred onto new RNAi plates before being tested for thermotolerance as previously described.

### Western blot analysis

For each strain, 1500 synchronized L1 worms were placed on NGM plates seeded with OP50 bacteria and incubated at 20°C for 48 h. These animals were then treated with Bortezomib (22 µM final) or control DMSO for 6 h and washed off the plate with M9. Proteins were extracted in lysis buffer (50 mM HEPES, pH7.4, 1 mM EGTA, 1 mM MgCl2, 100 mM KCl, 1% glycerol, 0.05% NP40, 0.5 mM DTT, 1x protease inhibitor tablet) as previously described (13). Protein levels were determined using the Pierce 660nm assay. Equal amount of proteins (5 µg) were boiled in protein loading buffer, separated on a 5-20% SDS-PAGE precast gel (Bio-Rad) and transferred onto PVDF membrane. Nonspecific binding was blocked using 5% nonfat dry milk in PBS-Tween (0.1%) for 1 hour at room temperature. The membranes were incubated with primary antibodies overnight at 4°C (mouse anti-FLAG diluted 1:1,000 and mouse anti-Tubulin diluted 1:7,500), washed 5 times in PBS-Tween and blotted in horseradish peroxidase-conjugated secondary antibodies at room temperature for 2 h (Goat anti-mouse diluted 1:10,000). Membranes were then washed 5 times in PBS-Tween, treated with ECL reagent (Amersham), and imaged using a Chemidoc XRS+ with Image Lab software (Bio-Rad).

### Fluorescence microscopy

For *cul-6, rcs-1* and *skr-5* tissue-specific expression lines shown in Fig. 1, Fig. 3 and Fig. 4, respectively, images were taken using a Zeiss LSM700 confocal microscope with 10X and 40X objectives. For the expression analysis of GFP::3xFLAG-tagged proteins PALS-22, PALS-25, CUL-6 and F42A10.5 in Fig S2, image were taken with a Zeiss LSM700 confocal microscope with 10X objective. For RNAi knock-down in Fig. S4, L4 stage F1 progeny of ERT422 worms fed with OP50 expressing *cul-6* RNAi were anesthetized using 10 µM levamisole in M9 buffer and mounted on 2% agarose pads for imaging with a Zeiss LSM700 confocal. Using ImageJ software (version 1.52e), GFP signal in the pharynx and the first intestinal cells ring was measured, as well as three adjacent background regions. The total corrected fluorescence (TCF) was calculated as TCF = integrated density – (area of selected cell × mean fluorescence of background readings). For each condition, 30 worms were imaged. Significance was assessed with a Student’s t-test.

### Measurement of developmental rate

40 – 50 gravid adults were transferred onto standard 10 cm NGM plate and incubated at 20°C to lay eggs for 2h before being removed. These plates were incubated at 20°C and the proportion of eggs that hatched and developed into L4’s was scored at 48 h, 64 h, and 72 h by scoring 100 animals each replicate.

## Acknowledgments

We thank Vladimir Lazetic, Ivana Sfarcic, Jessica Sowa, and Eillen Tecle for comments on the manuscript. This work was supported by NIH under R01 AG052622 to ERT and EJB, and a Burroughs Wellcome Fund Investigators in the Pathogenesis of Infectious Diseases fellowship to ERT.

## Supplementary Information

**Fig. S1.**
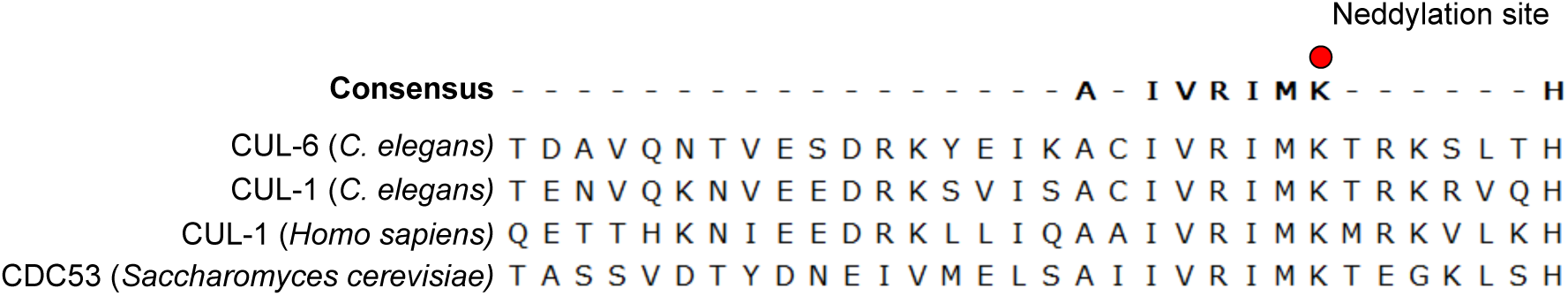
Alignment of the C-terminal region of *cul-6* with other cullin genes. The conserved lysine residue targeted by neddylation is indicated by a red circle.

**Fig. S2.**
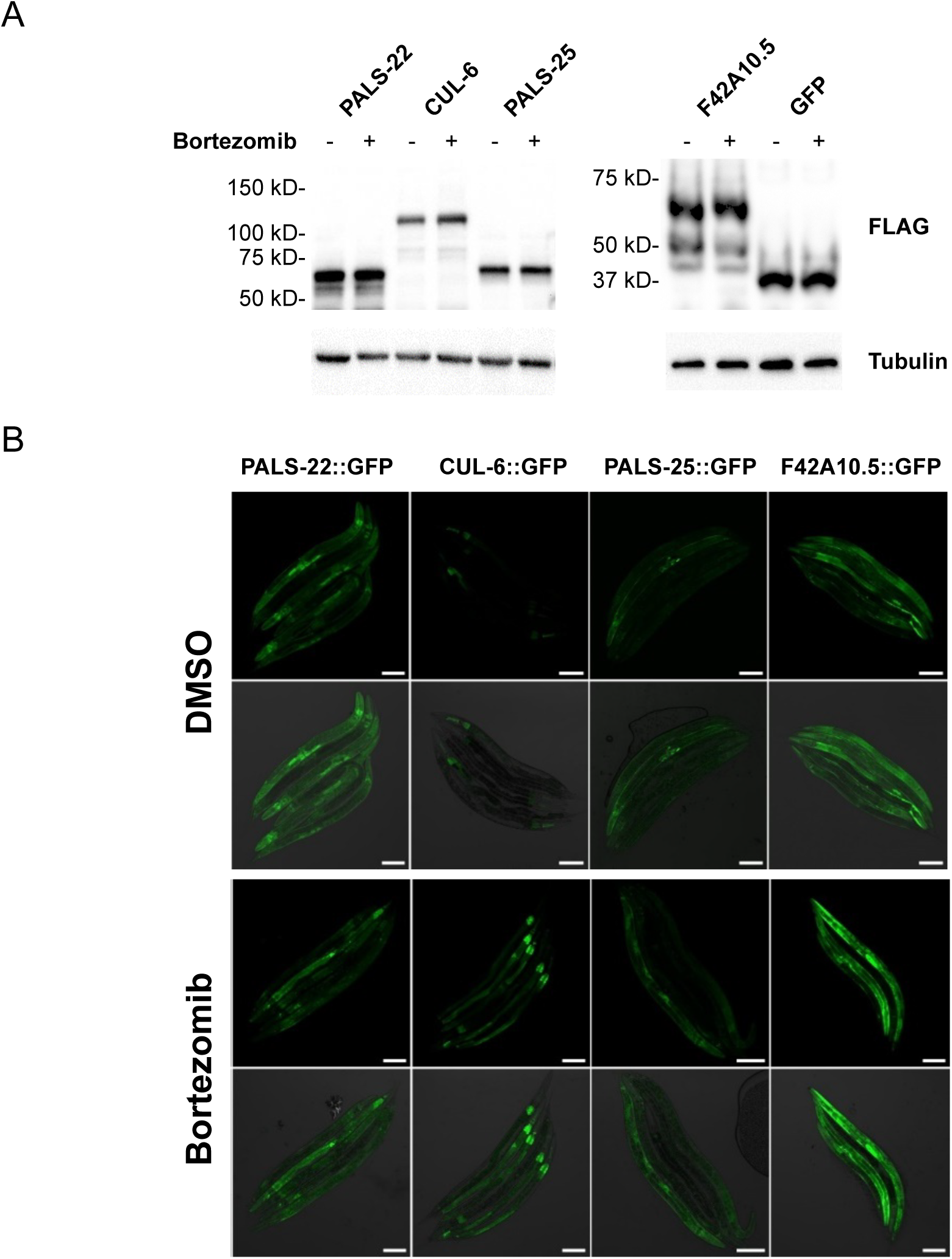
Expression analysis of GFP::3xFLAG-tagged proteins used for Co-IP/MS studies. *(A)* Western blot analysis of total protein lysate from transgenic adult animals containing fosmid transgenes expressing GFP::3xFLAG tagged protein and treated with Bortezomib or DMSO as diluent control. Proteins were detected with anti-FLAG, and anti-tubulin antibody was used as a loading control. Expected sizes; CUL-6::GFP::3xFLAG (116 kD), PALS-22::GFP::3xFLAG (64.8 kD), PALS-25 GFP::3xFLAG (66.9 kD), F42A10.5::GFP::3xFLAG (61.3 kD), GFP-3::FLAG (34 kD). *(B)* Confocal fluorescence images of L4 animals with fosmid transgenes expressing GFP tagged proteins from endogenous promoters, after exposure to DMSO or Bortezomib (diluted in DMSO). Images are overlays of green and phase contrast channels. Scale bar is 100 µm.

**Fig. S3.**
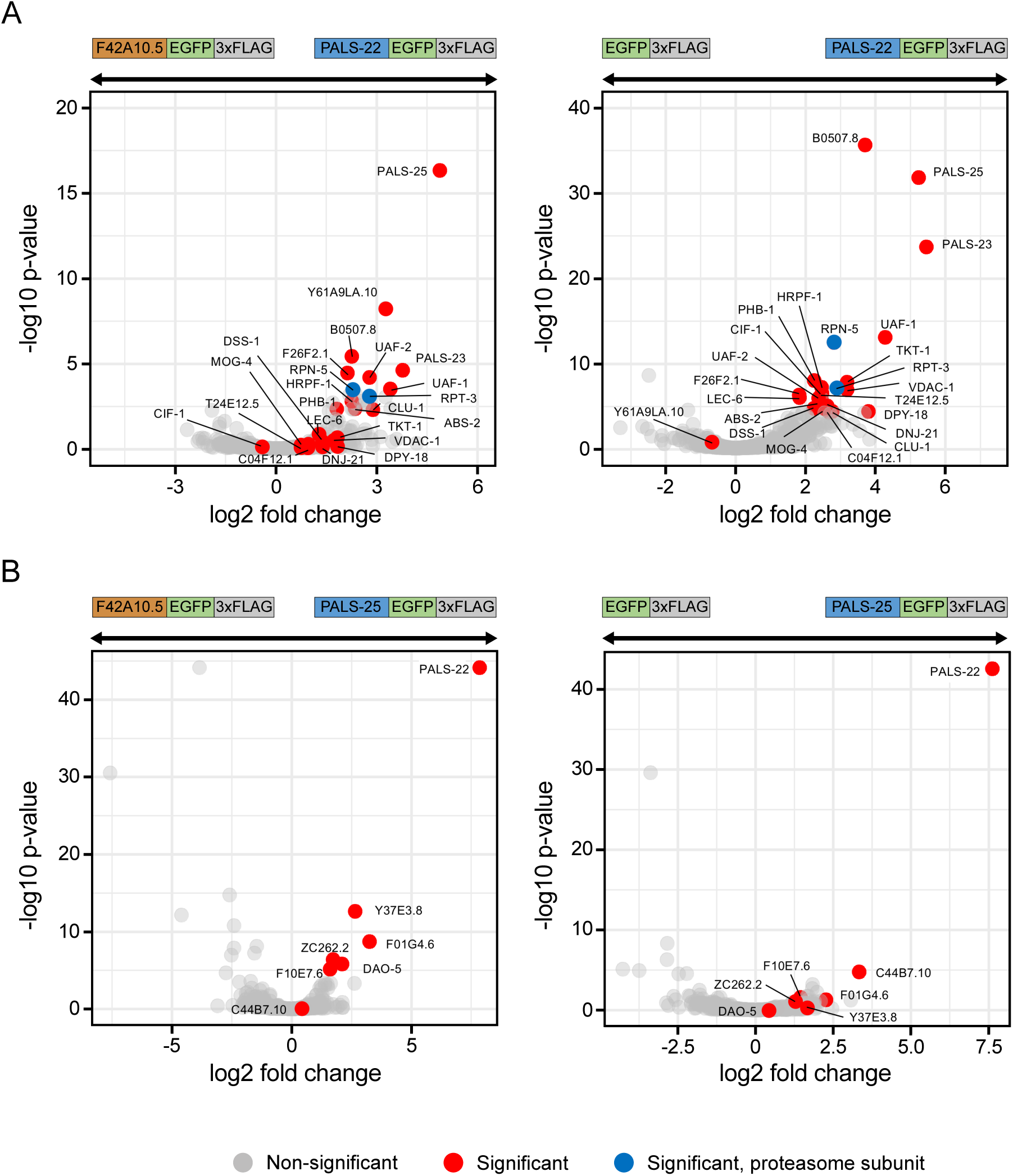
Co-immunoprecipitation mass spectrometry analysis identifies binding partners for PALS-22 and PALS-25. Volcano plot of proteins significantly enriched in PALS-22 *(A)* and PALS-25 *(B)* IP’s compared to F42A10.5 IP or GFP IP. Proteins significantly more abundant compared to either of the control IP’s (GFP alone control or F42A10.5 control, at adjusted *P* < 0.05 and log2 fold change > 1) were considered interacting proteins (Dataset S1). Gray dots indicate non-significant proteins, red dots indicate significant proteins and blue dots indicate significant proteasome subunits.

**Fig. S4.**
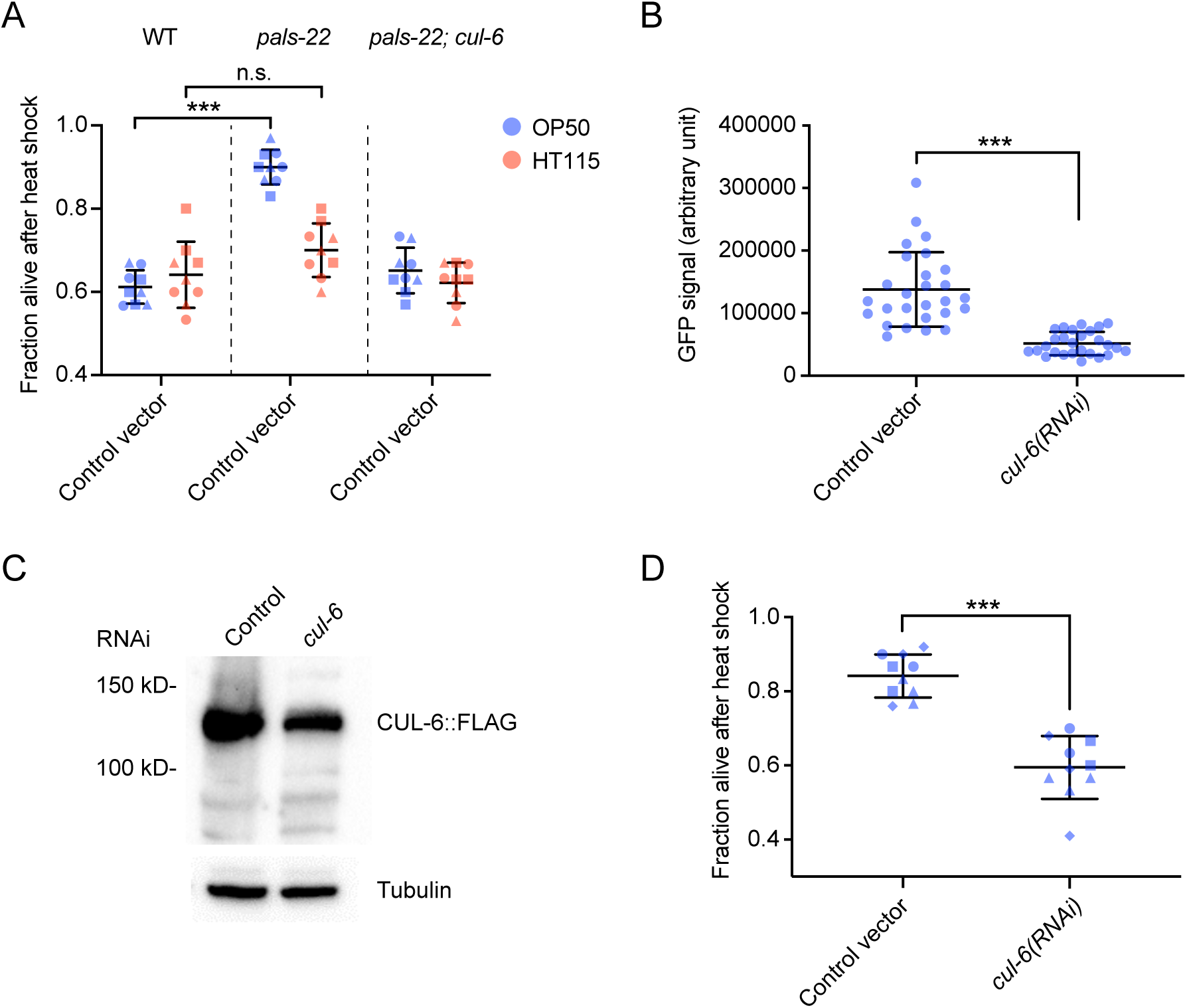
Development of an RNAi system for analyzing thermotolerance in *pals-22* mutants. *(A)* Survival of animals after 2 h of 37.5°C heat shock treatment, followed by 24 h at 20°C either fed on OP50 strain (R)OP50 or HT115 *E. coli*. Each dot represents a plate, and different shapes represent the experimental replicates done on different days. Mean fraction alive of the nine replicates is indicated by black bar with errors bars as SD. *** *P* < 0.001 with Student’s t-test. *(B)* Quantification of GFP signal in L4 animal expressing CUL-6::GFP grown on (R)OP50 expressing either L4440 (control vector) or *cul-6* RNAi. GFP Signal was measured with ImageJ in the pharynx and the first intestinal cells ring together with 3 adjacent background area and the Total Corrected Fluorescence (TCF) was calculated. Error bars are SD. *** *P* < 0.001 with Student’s t-test. *(C)* Western blot analysis on total protein lysate from adult animals with fosmid transgenes expressing CUL-6::GFP::3XFLAG treated with control or *cul-6* RNAi (OP50). CUL-6::GFP::3XFLAG protein was detected with anti-FLAG, and anti-tubulin antibody was used as a loading control. *(D)* Survival of *pals-22* mutants after 2 h of 37.5°C heat shock treatment, followed by 24 h at 20°C. Animals were fed on (R)OP50 expressing either L4440 (control vector) or *cul-6* RNAi. Each dot represents a plate, and different shapes represent the experimental replicates done on different days. Mean fraction alive of the nine replicates is indicated by black bar with errors bars as SD. *** *P* < 0.001 with Student’s t-test.

**Fig. S5.**
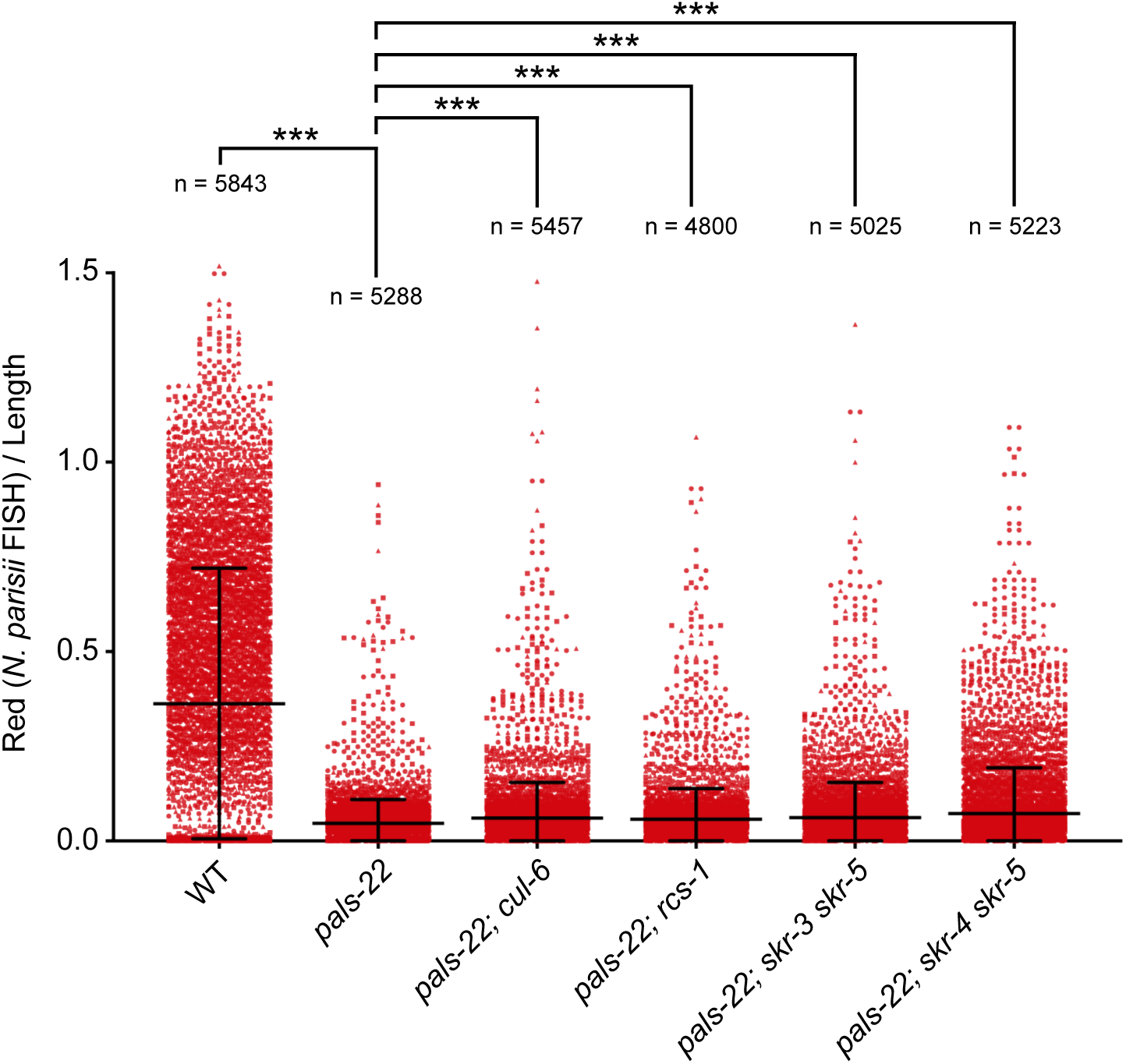
Analysis of pathogen resistance for *pals-22, cul-6, rcs-1* and *skr* double mutants in a *pals-22* mutant background. *N. parisii*-specific FISH probe (red) was quantified using a COPAS Biosort machine as mean red fluorescence normalized by length of individual animals. Strains were tested in triplicate infection experiments, three plates per experiment, approximately 1200 animals per plate. Each dot represents an individual animal and different shapes represent infection experiments conducted on different days. The number of animals analyzed across three separate infections is indicated for each strain. *** *P* < 0.001, Student’s t-test as compared to *pals-22* mutants. Error bars indicate mean and SD.

**Fig. S6.**
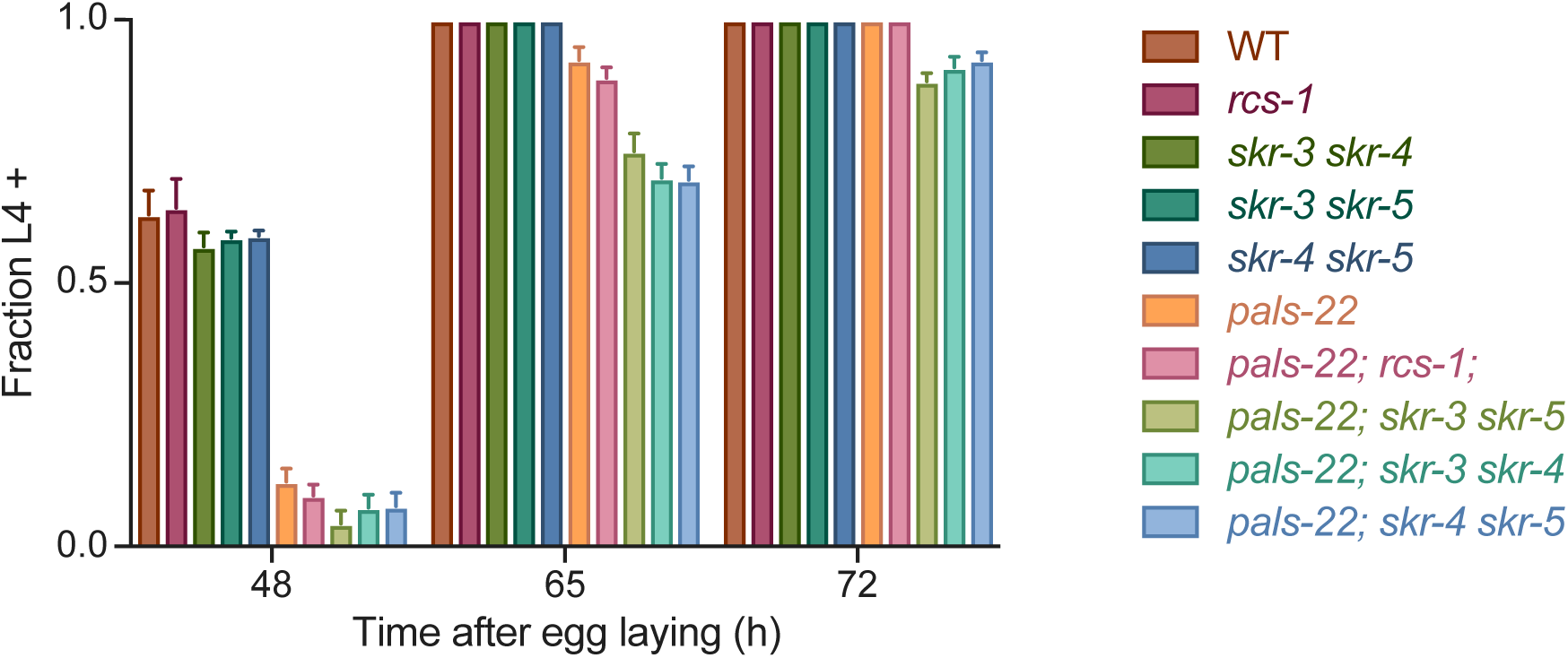
Analysis of developmental timing for *pals-22, rcs-1* and *skr* mutants. *skr* double and triple mutants in *pals-22* background do not suppress the developmental delay of *pals-22* mutants. Percentage of animals reaching the L4 larval stage at time points after eggs were laid is indicated. Results shown are the average of 3 independent biological replicates, with 100 animals assayed in each replicate.

**Table S1.** List of *C. elegans* strains used in this publication.

| Strain Name | Genotype (transgene or mutant allele details) | Source |
| --- | --- | --- |
| N2 | Wild type | Caenorhabditis Genetics Center |
| EG6699 | *ttTi5605 II; unc-119(ed3) III* | (Frøkjær-Jensen *et al.*, 2012) |
| RB2266 | *skr-5(ok3068) V* | Caenorhabditis Genetics Center |
| ERT54 | *jyls8[pals-5p::GFP; myo-2::mCherry] X* | (Bakowski *et al.*, 2014) |
| ERT356 | *pals-22(jy1) III* | (Reddy *et al.*, 2017) |
| ERT365 | *unc-119(ed3) III; jyEx193[pals-22::EGFP::3xFLAG, unc-119(+)]* | (Reddy *et al.*, 2017) |
| ERT413 | *jySi21[pET555(spp-5p::streplI3xFLAG GFP::let858 3'UTR; unc-119(+))] II; unc-119(ed3) III* | This paper |
| ERT422 | *unc-119(ed3) III; jyEx224[cul-6::EGFP::3xFLAG, unc-119(+)]* | (Reddy *et al.*, 2017) |
| ERT441 | *pals-22(jy1) III; cul-6(ok1614) IV* | (Reddy *et al.*, 2017) |
| ERT443 | *jySi22[pET592(myo-2p::GFP::APX NLS::unc-54; unc-119(+))] II; unc-119(ed3) III* | (Reddy *et al.*, 2017) |
| ERT465 | *unc-119(ed3) III; jyEx237[pals-25::EGFP::3xFLAG, unc-119(+)]* | (Reddy *et al.*, 2019) |
| ERT479 | *pals-22(jy3) III; skr-4(gk759439) V* | (Reddy *et al.*, 2017) |
| ERT488 | *unc-119(ed3) III; jyEx253[F42A10.5::EGFP::3xFLAG, unc-119(+)]* | (Reddy *et al.*, 2017) |
| ERT554 | *pals-22(jy1) III; skr-3(ok365) V* | (Reddy *et al.*, 2017) |
| ERT555 | *pals-22(jy1) III; skr-5(ok3068) V* | (Reddy *et al.*, 2017) |
| ERT571 | *jySi42[pET499(vha-6p::GFP::cul-6::unc-54 3' UTR, unc-119(+))] II; unc-119(ed3) III* | This paper |
| ERT638 | *jySi42 II; unc-119(ed3) pals-22(jy1) III; cul-6(ok1614) IV* | This paper |
| ERT691 | *rsc-1(jy84) X* | This paper |
| ERT692 | *pals-22(jy1) III; rcs-1(jy84) X* | This paper |
| ERT694 | *skr-3(ok365) skr-5(ok3068) V* | This paper |
| ERT717 | *skr-3(ok365) skr-4(gk759439) V* | This paper |
| ERT718 | *skr-5(ok3068) skr-4(gk759439) V* | This paper |
| ERT720 | *pals-22(jy1) III; skr-3(ok365) skr-4(gk759439) V* | This paper |
| ERT721 | *pals-22(jy1) III; skr-3(ok365) skr-5(ok3068) V* | This paper |
| ERT722 | *pals-22(jy1) III; cul-6(ok1614) IV; rcs-1(jy84) X* | This paper |
| ERT723 | *rsc-1(jy105) X* | This paper |
| ERT724 | *pals-22(jy1) III; rcs-1(jy105) X* | This paper |
| ERT727 | *pals-22(jy1) III; skr-5(ok3068) skr-4(gk759439) V* | This paper |
| ERT739 | *jySi45[pET686(myo-2p::GFP::cul-6::unc-54 3' UTR, unc-119(+))] II; unc-119(ed3) III* | This paper |
| ERT740 | *jySi46[pET688(vha-6p::GFP::cul-6(K673R)::unc-54 3' UTR, unc-119(+))] II; unc-119(ed3) III* | This paper |
| ERT741 | *jySi45 II; pals-22(jy1) unc-119(ed3) III; cul-6(ok1614) IV* | This paper |
| ERT742 | *pals-22(jy1) III; cul-6(ok1614) IV; rcs-1(jy105) X* | This paper |
| ERT752 | *jySi46 II; pals-22(jy1) unc-119(ed3) III; cul-6(ok1614) IV* | This paper |
| ERT817 | *jySi42 II; unc-119(ed3) III; skr-3 (ok365) skr-5 (ok3068) V* | This paper |
| ERT819 | *unc-119(ed3) III; jyEx286[pET711(rcs-1::EGFP::3xFLAG, unc-119(+))]* | This paper |
| ERT820 | *unc-119(ed3) III; jyEx287[pET561(skr-5::EGFP::3xFLAG, unc-119(+))]* | This paper |
| ERT852 | *fbxa-158(jy145) II* | This paper |
| ERT853 | *fbxa-158(jy146) II* | This paper |
| ERT854 | *fbxa-158(jy145) II; pals-22(jy1) III* | This paper |
| ERT855 | *fbxa-158(jy146) II; pals-22(jy1) III* | This paper |

**Table S2.** List of DNA constructs used in this publication.

| Construct Name | Genotype (transgene or mutant allele details) | Source |
| --- | --- | --- |
| pET499 | *vha-6p::SBP_3xFLAG_GFP_cul-6::unc-54 3'UTR* in pCFJ150 | This paper |
| pET555 | *spp-5p::streplI3xFLAG_GFP::let-858-3'UTR* in pCFJ150 | This paper |
| pET686 | *myo-2p::SBP_3xFLAG_GFP_cul-6::unc-54 3'UTR* in pCFJ150 | This paper |
| pET687 | *myo-3p::SBP_3xFLAG_GFP_cul-6::unc-54 3'UTR* in pCFJ150 | This paper |
| pET688 | *vha-6p::SBP_3xFLAG_GFP_cul-6(K673R)::unc-54 3'UTR* in pCFJ150 | This paper |
| pCFJ150 - Addgene Plasmid #19329 | *pDESTttTi5605[R4-R3]* | (Frøkjær-Jensen *et al.*, 2012) |
| pCFJ601 - Addgene Plasmid #34874 | *eft-3p::Mos1* transposase | (Frøkjær-Jensen *et al.*, 2012) |
| pMA122 - Addgene Plasmid #34873 | *peel-1* negative selection | (Frøkjær-Jensen *et al.*, 2012) |
| pGH8 - Addgene Plasmid #19359 | *rab-3p::mCherry* | (Frøkjær-Jensen *et al.*, 2012) |
| pCFJ90 - Addgene Plasmid #19327 | *myo-2p::mCherry* | (Frøkjær-Jensen *et al.*, 2012) |
| pCFJ104 - Addgene Plasmid #19328 | *myo-3p::mCherry* | (Frøkjær-Jensen *et al.*, 2012) |
| 873721959883807 G09 | *cul-6::EGFP::3xFLAG* Fosmid; *unc-119* selection | (Sarov *et al.*, 2012) |
| 9830596596427236 B12 | *pals-22::EGFP::3xFLAG* Fosmid; *unc-119* selection | (Sarov *et al.*, 2012) |
| 18995122782808704 G01 | *pals-25::EGFP::3xFLAG* Fosmid; *unc-119* selection | (Sarov *et al.*, 2012) |
| 8218370932910004 G03 | *F42A10.5::EGFP::3xFLAG* Fosmid; *unc-119* selection | (Sarov *et al.*, 2012) |
| 2491680425634929 B04 | *rsc-1::EGFP::3xFLAG* Fosmid; *unc-119* selection | (Sarov *et al.*, 2012) |
| 5202602939198744 E05 | *skr-5::EGFP::3xFLAG* Fosmid; *unc-119* selection | (Sarov *et al.*, 2012) |

### Dataset S1 (separate file)

Statistical analysis of co-IP experiments. Each tab shows the results for one experimental IP, CUL-6, PALS-22 or PALS-25. Each column indicates the fold change or adjusted p-value of the experimental IP relative to either F42A10.5 or GFP control IPs. Proteins indicated as “TRUE” were significantly more abundant in the experimental IP compared to control IPs (adjusted *P <* 0.05 and log2 fold change > 1).

